# Targeting conserved viral virulence determinants by single domain antibodies to block SARS-CoV2 infectivity

**DOI:** 10.1101/2021.01.13.426537

**Authors:** Sudhakar Singh, Surbhi Dahiya, Yuviana J. Singh, Komal Beeton, Ayush Jain, Roman Sarkar, Abhishek Dubey, Syed Azeez Tehseen, Sharvan Sehrawat

## Abstract

We selected SARS-CoV2 specific single domain antibodies (sdAbs) from a previously constructed phage display library using synthetic immunogenic peptides of the virus spike (S) protein as bait. The sdAbs targeting the cleavage site (CS) and the receptor binding domain (RBD) in S protein efficiently neutralised the infectivity of a pseudovirus expressing SARS-CoV2 S protein. Anti-CS sdAb blocked the virus infectivity by inhibiting proteolytic processing of SARS-CoV2 S protein. Both the sdAbs retained characteristic structure within the pH range of 2 to 12 and remained stable upto 65°C. Furthermore, structural disruptions induced by a high temperature in both the sdAbs were largely reversed upon their gradual cooling and the resulting products neutralised the reporter virus. Our results therefore suggest that targeting CS in addition to the RBD of S protein by sdAbs could serve as a viable option to reduce SARS-CoV2 infectivity and that proteolytic processing of the viral S protein is critical for infection.

## Introduction

The scale of infectivity and the spread of Severe Acute Respiratory SyndromeCoronavirus 2 (SARS-CoV2) caused COVID-19 pandemic continue to increase and the inflicted mortality is nearing two million worldwide^1^. There has been unprecedented progress in developing vaccines and therapeutics and some vaccines have been approved for use in some countries^2,3^. Many aspects of vaccine induced protective immunity such as the durability of response, the involvement of key immune mediators are yet to be adequately investigated. The development of potent yet affordable therapeutics such as anti-viral monoclonal antibodies is highly desirable not only to limit the consequences of COVID-19 but also to decipher events in virus entry as well as intracellular trafficking processes. Many such steps could provide useful anti-viral targets especially because the virus generates escape mutants that might be refractory to the induced immunity contributing to the enhanced virulence or transmissibility^4^. Passive immunotherapy with convalescent sera from recovered COVID-19 patients has limited utility and suffers from scalability issues^5^. Furthermore, the full-length conventional antibodies when administered in low abundance or those binding to the viral antigens with suboptimal avidities might contribute to antibody dependent enhancement (ADE) of infection^6,7,8^. Therefore, smaller variants of neutralizing antibodies such as fragment antigen binding (Fab), single chain fragment variable (scFv) or camelid single domain antibodies (sdAbs) could be considered as a promising approach^9,10^. Such interventions could reduce the viral loads to levels, which are efficiently controlled by immune mediators to restore homeostasis. Therefore, a potentially severe COVID-19 could be converted into a mild and manageable reaction. However, the commonly observed aggregation tendencies of Fabs and scFvs are yet to be satisfactorily resolved but the sdAbs are usually refractory to structural perturbations induced by biochemical and biophysical insults^11^. Such binders are well suited to neutralize infectious agents and toxins due to their ability to seek out cryptic epitopes^12^. Furthermore, SARS-CoV2 specific sdAbs could serve as valuable tool to investigate internalization, intracellular trafficking, replication, and assembly as well as the exocytic processes of the virus.

Membrane anchored spike (S) protein of SARS-CoV2 facilitates the virus entry in susceptible cells and is therefore considered as a major target for anti-viral manoeuvres^13,14^. Proteases such as furin, the host cell membrane associated TMPRSS2 process S protein by recognizing its polybasic residues (RRAR) in the cleavage site (CS) to generate S1 and S2 fragments^15,16^. This step is considered crucial for SARS-CoV2 entry, the viral fusion with endosomal membranes and to ensure the release of its genomes for translation and replication^17,18^. The acquisition of such a site by the newly emerged SARS-CoV2 is proposed to serve as a virulent factor for enhanced infectivity^19^. Not only SARS-CoV2 but many other viruses, bacterial products and toxins enhance their pathogenicity by such acquisitions and adaptations^20^. Antibodies targeting the RBD of SARS-CoV2 S protein have been selected and shown effective in the virus neutralization but those targeting the CS of SARS-CoV2 are not yet reported^21^. We therefore selected sdAbs against the CS and the RBD of S protein from a previously generated camelid **v**ariable region of **h**eavy chain of **h**eavy chain antibodies (V_H_H) phage display library^22^. We used synthetic peptides encompassing the cleavage site between S1/S2 as well as the linear epitopes present in the RBD of SARS-CoV2 S protein that were predicted to be immunogenic for selecting SARS-CoV2 specific sdAbs. Characteristically such sdAbs resisted structural disruptions and retained functionality even when exposed to harsh biophysical and biochemical conditions. We then demonstrated neutralizing potential of such sdAbs using a lentivirus based pseudovirus expressing surface SARS-CoV2 S protein as their entry mediator. Anti-CS sdAbs prevented the proteolytic cleavage of S protein and in so doing blocked the virus entry. Therefore, targeting multibasic CS as well as the RBD by sdAbs could be considered as a potential strategy to reduce SARS-CoV2 infectivity.

## Results and discussion

### Selection and characterization of SARS-CoV2 specific sdAbs

We predicted B cell epitopes of SARS-CoV2 S protein using the input amino acid sequence (PDB ID: 6VSB) described elsewhere^23^. The target regions of RBD were such selected that antibodies against such epitopes could interfere with the interactions involving the virus S protein and cellular entry receptors (Fig 1A). We also used a peptide encompassing the polybasic CS between S1/S2 fragments of S protein to select binders that could inhibit proteolytic processing by masking the site and in so doing inhibit the virus infectivity (Fig 1A, middle panel). Using synthetic peptides as bait, we biopanned specific sdAbs from a previously constructed phage display library without immunizing animals with SARS-CoV2 or its derivatives (Fig 1B,^24^). The sdAbs were expressed in *E. coli* (TG1 strain) and purified from the periplasmic fractions as well as inclusion bodies (Fig 1C and D, S1A-F). The purity and structural properties of the sdAbs were analysed by gel filtration, SDS-PAGE and circular dichroism (Fig 1C and D, and S1B-F). Fractions in peak 2 (P2) contained folded anti-CS sdAbs as revealed by a prominent peak at 218nm and a minor peak at 230nm in CD spectral plots, which indicated the presence of β sheets and aromatic amino acids in its secondary structure (Fig S1B, inset)^25^. Similarly, we selected, expressed and purified anti-RBD sdAb that were fully folded as revealed by CD plots (Fig 1D, S1D-F).

**Fig 1.**
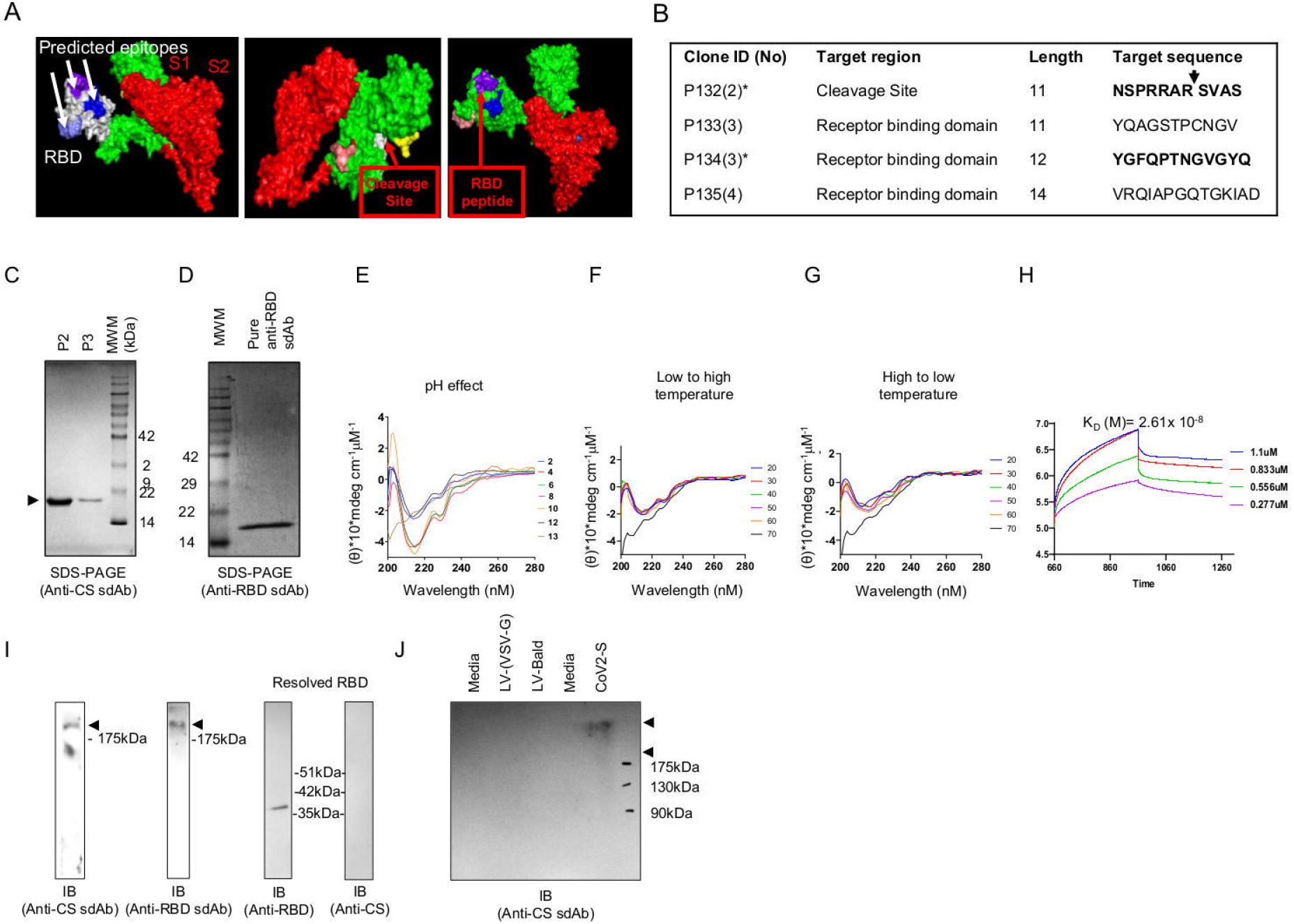
Generating SARS-CoV2 specific sdAbs by biopanning phage display libraries. A. Immunogenic peptides predicted for the SARS-CoV2 S protein are shown in the 3D structure. sdAbs against the boxed epitopes were selected and characterized. B. The clone ID, target region, numbers of residues in the peptides and sequences used for the selection of sdAb are tabulated. sdAb against the bold sequences were selected and characterised for measuring functionality. Downward arrow shows the site for cleavage in the peptide. C. Anti-CS sdAb was purified using Ni-NTA column and analysed by gel filtration chromatography. The resolved peaks (P2 and P3) using a 12% reducing SDS-PAGE were stained with CBBR-250. D. Anti-RBD sdAb was purified using Ni-NTA column and analysed using a 12% reducing SDS-PAGE. Stained gel image is shown. E-G. Anti-SARS-CoV2 sdAbs were characterised biophysically and biochemically. E. The influence of varying pH on secondary structure of anti-CS sdAb was measured by circular dichroism. The spectral analysis is shown for the anti-CS sdAb in indicated conditions. F. The influence of an increasing temperature from 20°C to 70°C on the secondary structure of anti-CS sdAb is shown by circular dichroism spectral analysis. G. The influence of decreasing the ambient temperature from 70°C to 20°C on the secondary structure of sdAb is shown by circular dichroism spectral analysis. H. The affinity of anti-CS sdAb against the SARS-CoV2 S protein preparation was determined by biolayer interferometry. The association and dissociation curve are shown and used to measure the affinity of anti-CS sdAb. I. The specificity of anti-CS and anti-RBD sdAb against the resolved recombinantly expressed S protein of SARS-CoV2 is shown by western blotting. J. The lysates from indicated samples; LV (VSV-G), LV (BALD) and SARS-CoV2 S transfected HEK293T cells were resolved using a 12% reducing SDS-PAGE followed by western blotting with anti-CS sdAb. All the experiments were repeated more than five times and representative images are shown.

We then analysed physicochemical robustness of sdAbs by exposing them to a range of pH and temperature (Fig 1E-G). The sdAbs retained their structural attributes between the pH ranges of 2 to 12 (Fig 1E). The sdAbs resisted structural perturbations when exposed upto a 65°C temperature (Fig 1F). Furthermore, the lost structures at high temperature (70°C) were regained with a gradual reduction in the storage temperature to 20°C (Fig 1G, data not shown). The affinity of anti-CS sdAbs with SARS-CoV2 S proteins was measured by biolayer interferometry and was found to be in the nanomolar range (Kd=2.6×10^-8^) (Fig 1H). The immune reactivity and the specificity of these sdAbs were determined against their selecting peptides as well as SARS-CoV2 S protein by western blotting and ELISA (Fig 1I and J, Fig 2). Anti-CS and anti-RBD sdAbs reacted with the virus S protein (~180kDa) while anti-RBD also reacted with recombinantly expressed RBD revealing a band of ~35kDa size (Fig 1I and J). The SARS-CoV2 S transfected HEK293T cells when probed with anti-CS sdAb revealed a specific band migrating above 180kDa molecular mass while no reactivity was observed against untransfected cells, assembled bald particles of pseudovirus, LV(BALD), or the VSV-G protein expressing pseudovirus, LV(VSV-G), resolved by a 12% SDS-PAGE followed by western blotting (Fig 1I and J).

**Fig 2.**
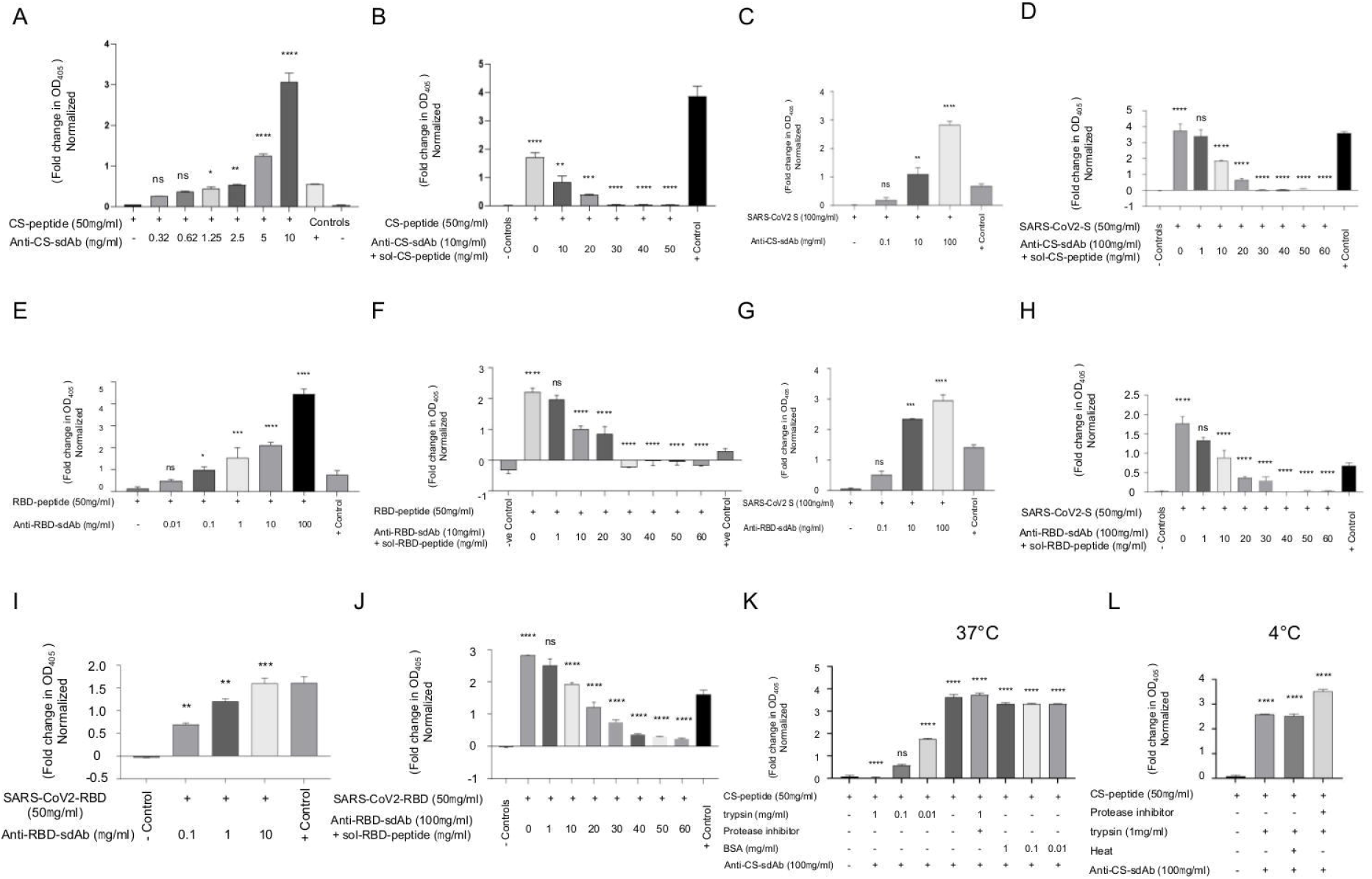
Determining the specificity and immune reactivity of anti-SARS-CoV2 S sdAbs. A. ELISA plates were coated with CS-peptide (50μg/ml) and probed using different concentrations of anti-CS sdAb. The fold change in the mean OD_405_ values as compared to mean OD_405_ values of negative controls is shown by bar diagram. B. ELISA plates were coated with CS-peptide (50μg/ml) and probed using anti-CS sdAb pre-incubated with the indicated concentrations of its specific peptide and a fold change in the OD_405_ values as compared to mean OD_405_ values of negative controls is shown by bar diagram. C. ELISA plates were coated with SARS-CoV2 S (100μg/ml) and probed with anti-CS sdAb. D. Anti-CS sdAb pre-incubated with different concentrations of soluble index peptide was used to probe immobilised SARS-CoV2 S protein and a fold change in the OD_405_ values in as compared to mean OD_405_ values of negative controls is shown by bar diagram. E. ELISA plates were coated with RBD-peptide (50μg/ml) and probed using different concentrations of anti-RBD sdAb and a fold change in the OD_405_ values as compared to mean OD_405_ values of negative controls is shown by bar diagram. F. ELISA plates were coated with RBD-peptide (50μg/ml) and probed using anti-RBD sdAb pre-incubated with the indicated concentrations of its specific peptide and a fold change in the OD_405_ values as compared to mean OD_405_ values of negative controls is shown by bar diagram. G. ELISA plates were coated with SARS-CoV2 S (100μg/ml) and probed using anti-RBD sdAb. H. Anti-RBD sdAb pre-incubated with different concentrations of soluble index peptide was used to probe immobilised SARS-CoV2 S protein and a fold change in the OD_405_ values as compared to mean OD_405_ values of negative controls is shown by bar diagram. I. ELISA plates were coated with recombinant RBD of SARS-CoV2 (50μg/ml) and probed using graded concentrations of anti-RBD sdAb and a fold change in the OD_405_ values as compared to mean OD_405_ values of negative controls is shown by bar diagram. J. ELISA plates were coated with CS-peptide (50μg/ml) and probed using anti-RBD sdAb pre-incubated with the indicated concentrations of its specific peptide and a fold change in the OD_405_ values as compared to mean OD_405_ values of negative controls is shown by bar diagram. K. ELISA plates were coated with 50μg/ml of CS-peptide alone or that previously incubated with different concentrations of trypsin for three hours at 37°C. In additional wells, the mix of CS-peptide was first incubated with protease inhibitors followed by addition of trypsin for three hours at 37°C. Some wells were additionally coated with the CS-peptide along with BSA under similar conditions. The plates were then probed using anti-CS sdAb and a fold change in the OD_405_ values as compared to mean OD_405_ values of negative controls is shown by bar diagram. L. ELISA plates were coated with 50μg/ml of CS-peptide alone or that previously incubated with different concentrations of trypsin for three hours at 4°C. In additional wells, the mix of CS-peptide was first incubated with protease inhibitors followed by addition of trypsin for three hours at 4°C. Some wells were additionally coated with the mix of CS-peptide and trypsin under similar conditions followed by its heating. The plates were then probed using anti-CS sdAb and a fold change in the OD_405_ values as compared to mean OD_405_ values of negative controls is shown by bar diagram. The experiments were repeated three times and representative data plots from one such experiment in the respective section is shown. One way ANOVA was used to measure the level of statistical significance. ****p<0.0001, ***p<0.001, *p<0.01 and *p<0.05.

The CS and the RBD peptides when probed with their specific sdAbs expectedly showed a concentration dependent increase in ELISA readouts (OD_405_) but flipping the probing sdAbs failed to do so (Fig 2A and E, S1G and H). Furthermore, a prior incubation of both the sdAbs separately with their selecting peptides in solution followed by probing against plate bound index peptides reduced signal intensity as the concentration of respective peptides increased (Fig 2B and F). These results showed specificity of sdAbs for their cognate peptides. We also demonstrated the concentration dependant reactivity of both the sdAbs against SARS-CoV2 S protein in ELISA (Fig 2C and G). Moreover, their prior incubation with index peptides inhibited the binding to immobilized SARS-CoV2 S protein albeit the response was less evident for anti-RBD sdAb (Fig 1D and H). A 30μg/ml of the CS peptide and a 40μg/ml of RBD peptides pre-incubated with their respective sdAbs completely erased the signal (Fig 2D and H). These results established the specific binding of anti-CS and anti-RBD sdAbs. We also probed anti-RBD sdAb against the recombinantly expressed RBD using ELISA and observed a concentration dependent increase in OD_405_ values that were reduced by prior incubation with the RBD peptide (Fig 2I and J)

We then attempted to map the binding sites of anti-CS sdAb in order to reveal its potential anti-viral mechanisms. Trypsin, a serine protease that recognises and cleaves basic residues, was incubated with the CS-peptide in varying concentrations at 37°C and the mix was coated to ELISA plates followed by its probing with anti-CS sdAb in ELISA (Fig 2K and L and Fig S1I). The signal intensity was gradually reduced as the concentration of trypsin increased (Fig 2K). Equivalent concentrations of a non-specific protein, bovine serum albumin, incubated with the peptides under similar conditions did not affect the assay readouts (Fig 2K). Furthermore, a prior incubation of the CS-peptide with protease inhibitor reversed these effects (Fig 2K). These results suggested that the proteolytic activity of trypsin could have erased the epitope recognised by anti-CS sdAb and in so doing diminished the response. We then determined whether these results could also be due to masking of epitope by trypsin and thereby making it unavailable for binding to the antibody. To this end, we incubated the peptide and trypsin mix at low temperature (4°C), a procedure that would minimize the proteolytic activity of trypsin but leaving the binding unaffected. The mix of CS-peptide and trypsin was additionally heat inactivated at 90°C to inhibit residual proteolytic activity before coating onto the plates. The increase in OD_405_ values in such assays indicated the crucial role of enzymatic activity in diminishing the signal (Fig 2L). We obtained essentially similar results when the SARS-CoV2 S protein was used (data not shown). These results could indicate that anti-CS sdAb recognized the epitope bordering S1/S2 segments in the unprocessed S protein of the SARS-CoV2. By extrapolation these experiments could suggest that anti-CS sdAb could make the CS inaccessible for proteolytic activity, a step that precedes SARS-CoV2 internalization.

Taken together, we selected SARS-CoV2 S protein specific sdAbs from a phage display library using synthetic peptides as the bait. We then established the robustness, thermostability, specificity and immune reactivity of SARS-CoV2 S protein specific sdAbs not only against their selecting peptides but also using SARS-CoV2 S protein.

### Anti-SARS-CoV2 sdAbs neutralize the virus infectivity and inhibit S protein mediated cell to cell fusion

Having established the biophysical and biochemical characteristics of SARS-CoV2 specific sdAbs, we measured their neutralizing activity against lentivirus (LV) based reporter pseudoviruses that express either SARS-CoV2 S or VSV-G protein (Fig 3). The use of pseudovirus for analysing neutralizing antibodies or other anti-viral agents could obviate the requirement of high containment facilities and the results from such assays correlate with neutralization of the virulent virus *ex vivo* or *in vivo*^26^. Furthermore, such a system can efficiently be used for testing the neutralization of emerging mutants without necessarily isolating the virulent viruses.

**Fig 3.**
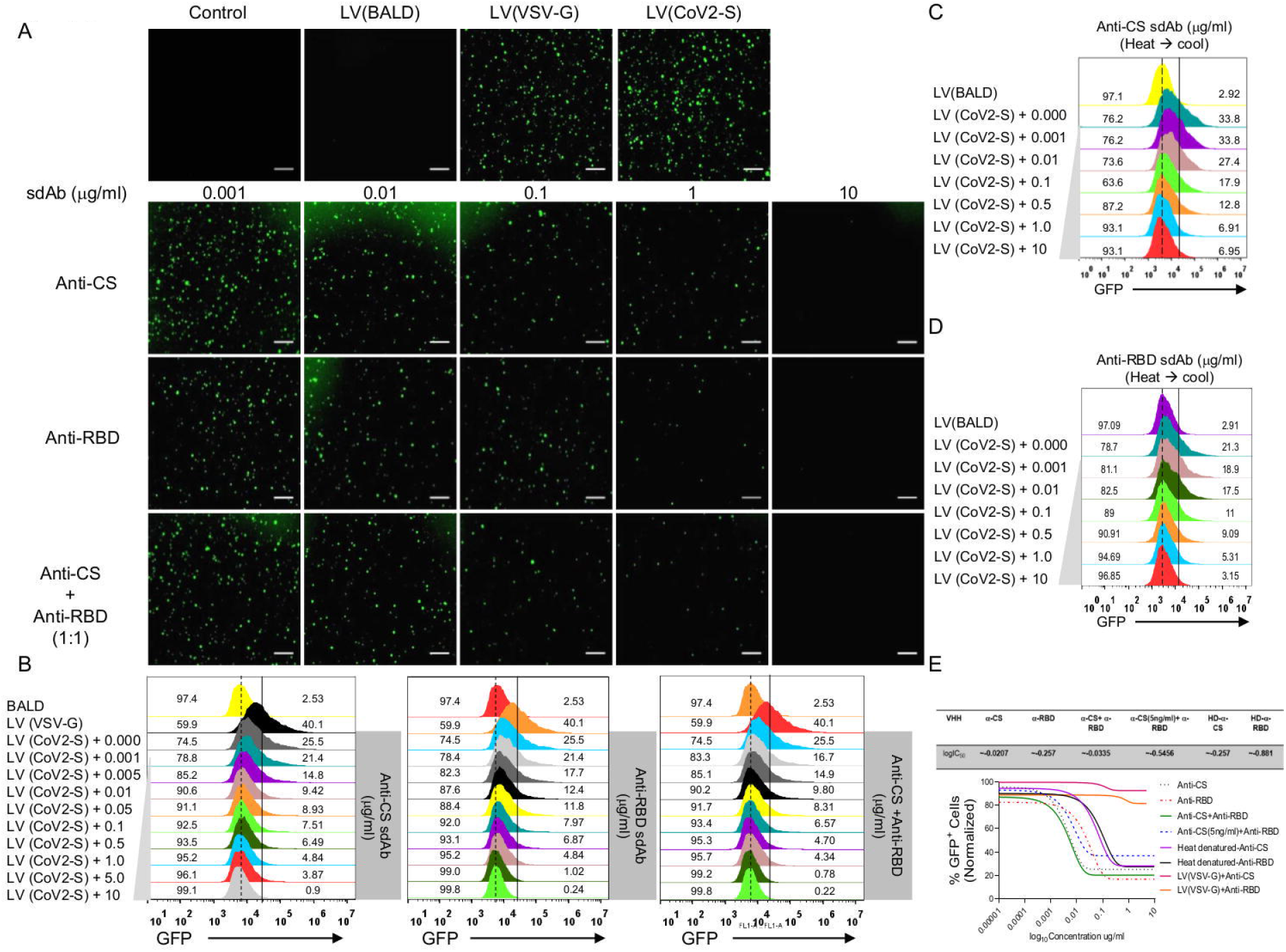
Anti-SARS-CoV2 S protein specific sdAbs neutralize the reporter virus. The neutralizing potential of Anti-SARS-CoV2 S protein specific sdAbs was analysed using lentivirus (LV) based pseudovirus expressing SARS-CoV2 S, LV (CoV2-S) and VSV-G, LV (VSV-G). Vero E6 cells were infected with the pseudovirus particles alone or pseudovirus particles pre-incubated with anti-CS and/or anti-RBD sdAbs at 4°C. Green fluorescence emitted by such cells was analysed by fluorescent microscopy and flow cytometry. A. Representative fluorescent microscopic images show the extent of neutralization of LV (CoV2-S) by different concentrations of anti-CS and/or anti-RBD sdAbs. For a measuring the combinatorial effect each of the sdAbs half of the doses were used, scale bar 160μm. B. Representative overlaid histograms obtained by flow cytometry show GFP^+ve^ cells infected with native LV (CoV2-S) infected Vero E6 cells or the anti-CS and/anti-RBD sdAb neutralized LV (CoV2-S) in indicated conditions. C. Representative overlaid histograms obtained by flow cytometry show GFP^+ve^ cells infected with native LV (CoV2-S) infected Vero E6 cells or the pre-incubated LV (CoV2-S) with high temperature exposed anti-CS sdAb followed by its cooling at room temperature. D. Representative overlaid histograms obtained by flow cytometry show GFP^+ve^ cells infected with native LV (CoV2-S) infected Vero E6 cells or the pre-incubated LV (CoV2-S) with high temperature exposed anti-RBD sdAb followed by its cooling at room temperature. Vertical dotted line serves as the reference for marking the peak and vertical dark line represents the marker to show GFP^+ve^ and GFP^-ve^ cells. E. Cumulative data obtained from neutralization experiments along with the inhibitory concentrations are depicted. The experiments were performed for more than five times with similar results. One way ANOVA was used to measure the level of statistical significance. ****p<0.0001, ***p<0.001, *p<0.01 and *p<0.05.

The scanning electron micrographs showed the presence of spikes on LV(CoV2-S) pseudovirus particles that were conspicuously missing by a control LV(VSV-G and such particles were 50nm-100nm in diameter (Fig S2A). The susceptible Vero E6 cells incubated with LV(CoV2-S) fluoresced within 24hrs and the percent positivity increased upto 72 hrs post incubation. We obtained upto a 50% GFP^+ve^ cells in different experiments (Fig 3A, B and E). A prior incubation of LV(CoV2-S) with varying concentration of anti-CS and/or anti-RBD sdAbs reduced cellular infectivity as revealed by fluorescence microscopy and flow cytometry (Fig 3A, B and E). A significant inhibition of the virus entry was observed at concentrations of anti-CS and/or anti-RBD sdAbs as low as 1ng/ml (Fig 3A, B and E, Fig S2B). Upto a 50% neutralization was achieved at 100ng/ml by both the sdAbs. At 5μg/ml of either the sdAbs achieved a near complete inhibition of the virus infectivity (Fig 3A, B and E). The inhibitory concentration (IC_50_) values were ~10 times lower for anti-CS sdAb as compared to anti-RBD sdAb (Fig 3E). We also measured whether or not combining both the sdAbs enhanced the efficiency of neutralization and observed a slight improvement in the neutralization efficiencies particularly at lower concentrations when both the sdAbs (each with 0.5ng/ml) were added in similar assays (Fig 3C). We also observed an improved neutralization when a fixed but sub-optimal concentration (5ng/ml) of anti-CS sdAb was combined with varying concentrations of anti-RBD sdAb but a general potentiation effect was less evident (Fig 3A, B, E and Fig S3). We also assessed the specificity of blockade by both the sdAbs using LV(VSV-G) or by their prior incubation with the cognate peptides (Fig 3E, Fig S2C-F and Fig S4). None of the sdAbs prevented infectivity of LV(VSV-G) even in ~10,000 molar excess values as were used for neutralizing LV(CoV2-S) (Fig 3E and Fig S2C-F). Furthermore, a prior incubation of both the sdAbs with increasing concentrations of their cognate peptides significantly reduced the virus neutralization efficiencies (Fig S4). These results clearly demonstrated the specificity of virus neutralization by both the sdAbs through recognition of their cognate viral epitopes displayed by SARS-CoV2 S protein in LV(CoV2-S). The heat denatured sdAbs regained structural features as the ambient temperature was reduced (Fig 1G). Anti-CS and anti-RBD sdAbs upon their renaturation neutralized the virus infectivity albeit with a reduced efficiency (Fig 3C-E). Accordingly, the IC50 concentrations for renatured anti-CS sdAb and anti-RBD sdAbs were respectively ~10 and 4 fold lower in comparisons to their native preparations (Fig 3E). These results demonstrated the neutralization of a reporter pseudovirus expressing SARS-CoV2 S protein by the sdAbs, which also withstood harsh biochemical and biophysical conditions. Furthermore, their lost structural and functional properties induced by a high temperature were largely regained with lowering of the storage temperature attesting to their enhanced utility.

SARS-CoV2 causes cell to cell fusion using its surface exposed S protein and in so doing infects bystander cells^27^. We therefore tested whether or not anti-CS and anti-RBD sdAbs could block fusogenic activity. HEK293T cells were used for producing LV(CoV2-S) pseudovirus particles, which could remain cell associated during the exocytic process. The synthesized S proteins might be displayed by infected cells. HEK293T cells used for producing SARS-CoV2 expressing pseudoviruses (HEK293T^+LV(CoV2-S)^) or the control particles (HEK293T^+LV(BALD)^) were co-cultured with Vero E6 cells in the presence or absence of sdAbs (Fig 4A-C). HEK293T^+LV(BALD)^ did not fuse with Vero E6 cells as revealed by clear GFP punctates (Fig 4A, upper middle panel). HEK293T^+LV(CoV2-S)^ induced high fusogenic activity with Vero E6 cells as revealed by extensively diffused pattern of GFP fluorescence (Fig 4A, upper right panel, Fig S5). A pre-incubation of HEK293T^+LV (CoV2-S)^ cells with the graded concentrations of anti-CS sdAb, anti-RBD sdAb or a combination thereof reduced the fusogenic activity by upto five-fold (Fig 4A-C). Accordingly, we observed ~25% GFP^+ve^ cells in the absence and ~5% GFP^+ve^ cells in the presence of either of the sdAb preparation (Fig 4B-C). Similarly, both the sdAbs either alone or in combination reduced the fusogenic activity by four-fold when transfected HEK293T cells (HEK293T^+(CoV2-S)^) were incubated with Vero E6 cells (Fig 4D-F). For such fusion events to occur, the expression of the viral S protein on cell surface is a prerequisite^28^. We therefore measured the surface expression of S protein in HEK293T cells by flow cytometry using both the sdAbs (Fig 4G). We used biotinylated anti-CS and anti-RBD sdAbs to detect expressed S protein by HEK293T cells. While the control cells (HEK293T^+LV(BALD)^) showed no staining with either of the sdAbs, the staining for the S protein was clearly evident in the transfected cells. We observed >40% and >65% CoV2-S positive cells detected by anti-CS sdAb and anti-RBD sdAb, respectively (Fig 4G).

**Fig 4.**
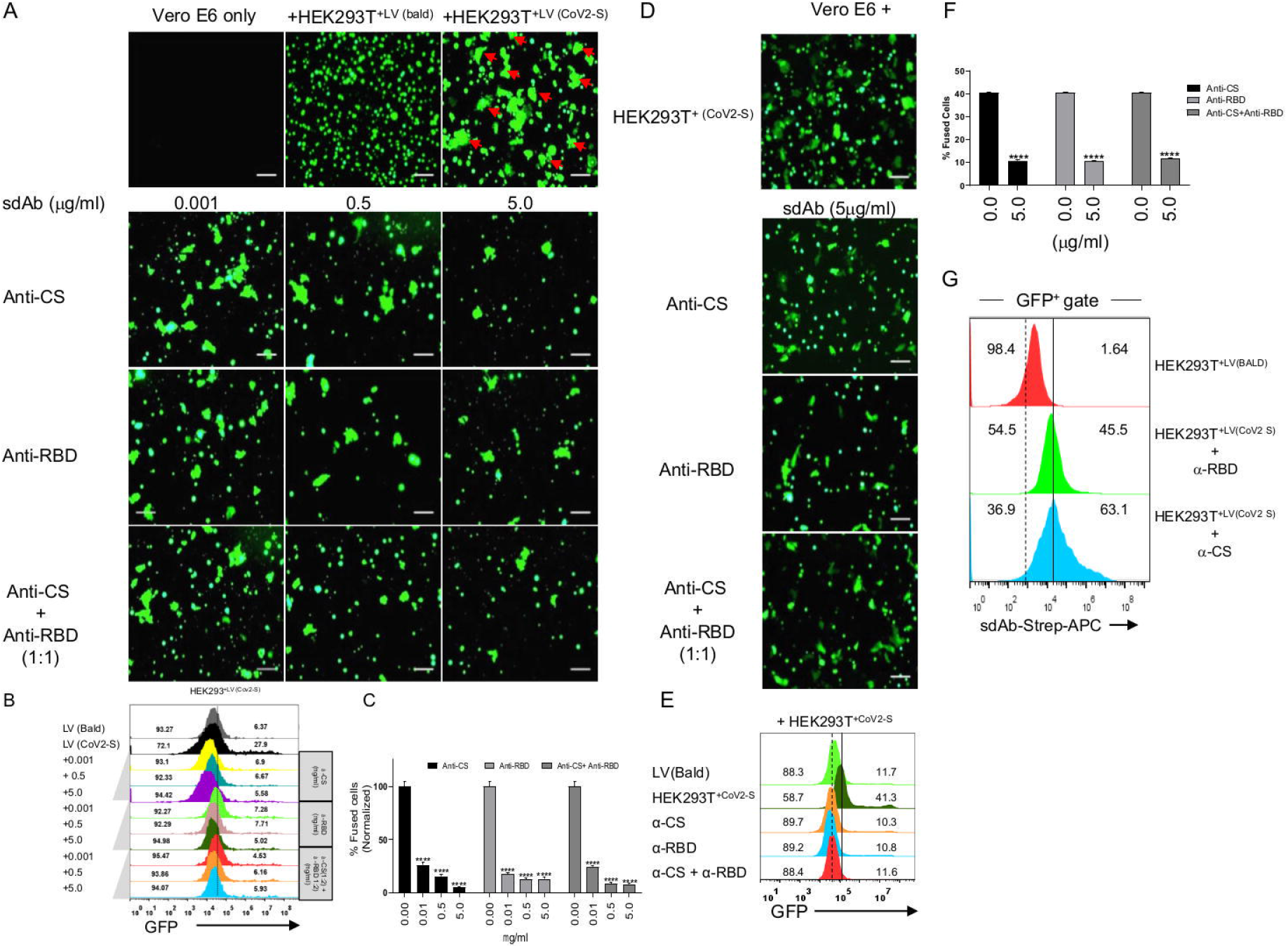
Anti-SARS-CoV2 S protein sdAbs inhibit cell to cell fusion. HEK293T cells used for producing LV(CoV2-S), HEK293T^+LV(CoV2-S)^, and LV(BALD), HEK293T^+LV(BALD)^ were cocultured with Vero E6 cells in the presence or absence of anti-CS and/or anti-RBD sdAb and the percentage of fused cells was quantified by fluorescent microscopy and flow cytometry. A. Representative fluorescent images from indicated conditions are shown. Red arrow heads indicate fused cells (diffused GFP staining) in the co-culture of Vero E6 cells and HEK293T^+LV(CoV2-S)^ cells while no such staining pattern was observed in the co-culture of Vero E6 cells and HEK293T^+LV(BALD)^ cells. The extent of fused cells is shown when HEK293T^+LV (CoV2-S)^ cells were pre-incubated with anti-CS and/or anti-RBD sdAb when used alone or in combination at indicated concentrations, scale bar 160 μm. B. Representative overlaid histograms show the percentage of fused cells in different conditions. Gating strategy for determining the percentage of fused cells is shown in Fig S5. C. Bar diagrams show the percentage of normalized fused cells in co-culture with or without different concentrations of anti-CS or/and anti-RBD sdAbs. D. The fusogenic response of HEK293T cells transfected with SARS-CoV2 S construct, HEK293T^(CoV2-S)^ when cultured with Vero E6 cells in the presence or absence of anti-CS and/or anti-RBD sdAbs is shown by fluorescent images scale bar 160μm. E. Representative overlaid histograms show the percentage of fused cells in different conditions. Gating strategy for determining the percent fused cells is shown in Fig S5. F. Bar diagrams depict the percentage of fused cells in co-culture with or without different concentrations of anti-CS or/and anti-RBD sdAbs. G. Surface expression of SARS-CoV2 S protein by HEK293T^+LV(CoV2-S) was^ measured by flow cytometry using biotinylated anti-CS and anti-RBD sdAbs and representative histograms show percent positive cells. The experiments were repeated four times with similar results. The levels of statistical significance were measured by one way ANOVA. ****p<0.0001, ***p<0.001, *p<0.01 and *p<0.05.

Taken together our results showed that SARS-CoV2 S specific sdAbs block not only the virus entry in susceptible cells but also prevent its fusogenic activity.

### Anti-CS sdAb inhibits SARS-CoV2 infectivity by preventing proteolytic cleavage of the S protein

We observed that a prior incubation of CS-peptide with trypsin reduced the assay readouts (Fig 2K, L and Fig S1I). These observations led us to explore possible mechanisms by which anti-CS sdAb functions to neutralise the virus (Fig 5). We probed the culture supernatants from Vero E6 cells infected with either control LV(CoV2-S) or anti-CS sdAb treated LV(CoV2-S) with an anti-FLAG antibody. An intense band migrating at ~180kDa molecular mass corresponding to the unprocessed spike protein (S0) was observed and its intensity increased in samples where increasing concentration of anti-CS sdAb was added (Fig 5A, Fig S6A-C). This suggested for an abundance of LV(CoV2-S) particles in culture supernatants due to less efficient internalisation upon anti-CS antibody addition. We then tested whether or not anti-CS sdAbs inhibited the cleavage of S protein. LV(CoV2-S) were incubated with either anti-CS or anti-RBD sdAbs followed by a trypsin treatment. Four hours later the mix were analysed by immunoblotting using anti-FLAG antibodies. A pre-incubation of LV(CoV2-S) with anti-CS sdAb but not with anti-RBD sdAb prevented the processing of S protein as revealed by intense band of ~180kDa (Fig 5B). In other condition the band of ~180kDa was barely detected (Fig 5B). A prior incubation of LV(CoV2-S) with either of the sdAbs followed by trypsin treatment significantly inhibited the internalization process nonetheless. This observation suggested that both the antibodies recognised distinct epitopes in the S protein and acted independently to inhibit the virus internalization (Fig 5C, Fig S7A). We also pre-treated LV(CoV2-S) with different concentrations of trypsin followed by incubation with the anti-CS sdAb (Fig 5D-F). In these experiments, the anti-CS sdAb (100ng/ml) significantly neutralized native LV(CoV2-S) by ~50% but the trypsin treated LV(CoV2-S) particles remained refractory to the antibody (Fig 5D-F). These results suggested that the anti-CS sdAb blocked SARS-CoV2 infectivity by recognizing the unprocessed S protein (S0) displayed by LV(CoV2-S) and prevented its proteolytic processing. Therefore, the processing of S protein is a crucial step in viral entry. We also tested whether anti-RBD sdAb by binding to a different site in the S protein could inhibit the virus internalization by independently of its proteolytic processing. A prior trypsin treatment of LV(CoV2-S) followed by its incubation with the graded concentrations of anti-RBD sdAb efficiently blocked the virus entry (Fig S7B-D). Moreover, coated LV(CoV2-S) on the surface of plates in native form showed reactivity with anti-CS sdAb but its prior treatment with trypsin reduced the signal intensity in a dose dependent manner (Fig S7E). The reactivity remained unchanged with anti-RBD sdAb (Fig S7F). These results established not only the specificity of sdAbs to recognise epitopes displayed by S protein in the assembled LV(CoV2-S) but also hinted for the functional diversity of both the sdAbs in effecting the neutralization the pseudovirus particles.

**Figure 5.**
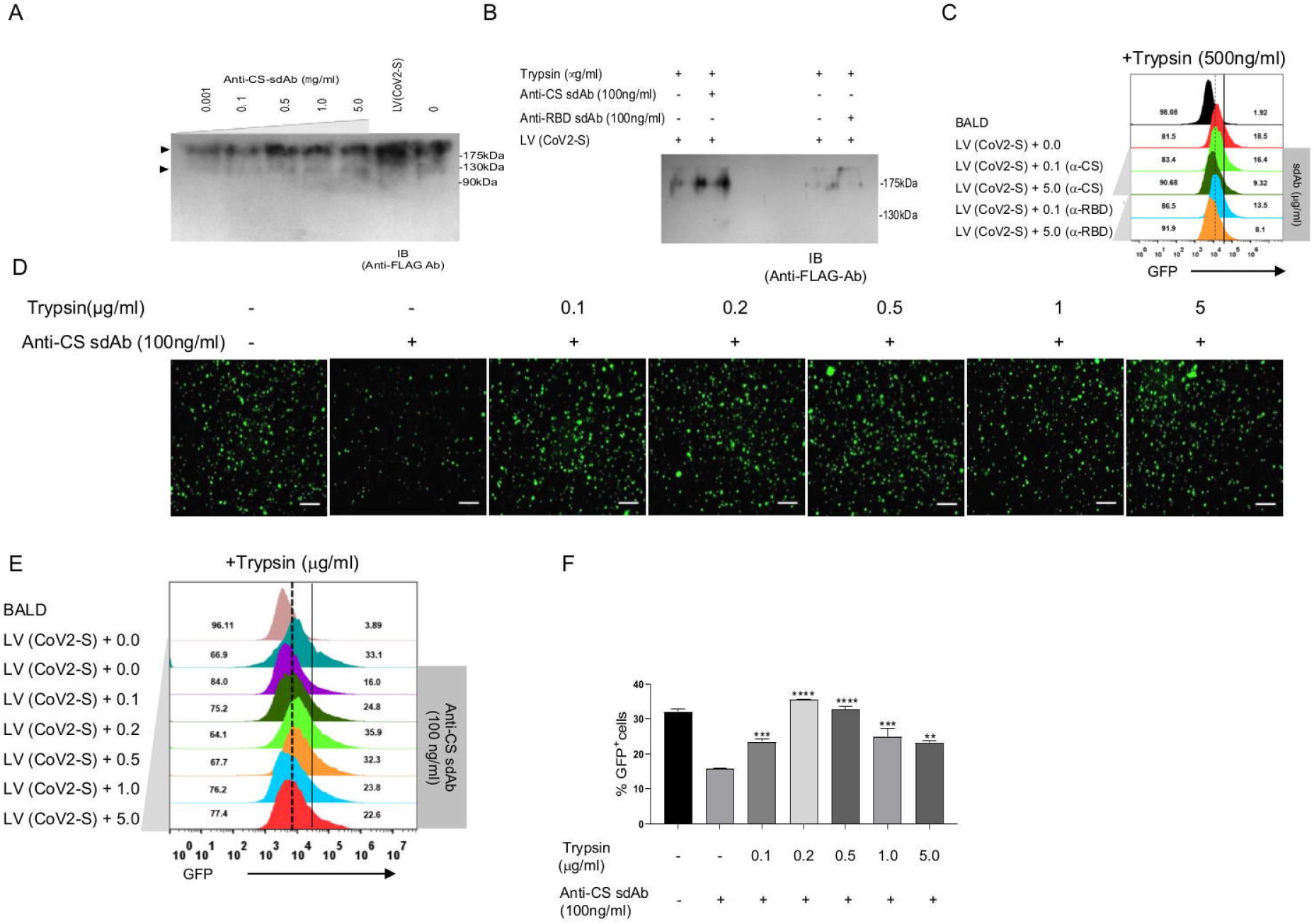
Deciphering possible anti-viral mechanism of anti-CS sdAb. A. Vero E6 cells were infected with native LV (CoV2-S) or those pre-incubated with varying concentrations of anti-CS sdAbs. After 72 hour post incubation, culture supernatants were collected from each condition and after processing were resolved by a 12% reducing SDS-PAGE followed by transferring polypeptides to PVDF membranes. The membranes were then probed with anti-FLAG monoclonal antibody. Immunoblot show the presence of polypeptide bands ~180kDa and ~95kDa range to corresponding to full length unprocessed S protein (S0) and cleaved product (S2) indicated by upper and lower arrowheads, respectively. B. LV (CoV2-S) was incubated with 5μg/ml of either anti-CS or anti-RBD sdAb followed by its treatment with 500ng/ml of trypsin for 4 hrs at 37°C. The cocktail was resolved by a 12% reducing SDS-PAGE followed by transfer to PVDF membrane. The membranes were then probed using anti-FLAG monoclonal antibody. More of the unprocessed protein (S0) was observed when anti-CS sdAb was added before trypsin treatment. C. LV (CoV2-S) processed in B. were added to Vero E6 cells and the percentage of GFP^+ve^ cells were analysed by fluorescent microscopy (Fig S7) and flow cytometry. Representative overlaid histograms show GFP^+ve^ and negative cells in different conditions. D. The fluorescent microscopic images show the neutralization of native or trypsin treated LV-(CoV2-S) infectivity in Vero E6 cells by anti-CS sdAb scale bar 160μm. E. Overlaid histogram obtained by flow cytometry show GFP^+ve^ and negative Vero E6 cells in indicated conditions. F. Bar diagram show cumulative data to assess the neutralizing effects of anti-CS sdAb against LV-(CoV2-S) in Vero E6 cells. All the experiments were repeated three times and representative images and flow cytometric histograms are shown. One way ANOVA was used to measure the level of statistical significance. ****p<0.0001, ***p<0.001, *p<0.01 and *p<0.05.

## Conclusion

We describe here neutralizing sdAbs selected from a phage display library of camelid VHH that target RBD and CS of SARS-CoV2 S protein. While the neutralizing anti-RBD sdAbs have been reported, anti-CS sdAb targeting the CS of SARS-CoV2 have not been selected^29^. Furthermore, as a point of departure we used synthetic immunogenic peptides of SARS-CoV2 S protein for efficiently selecting the virus neutralizing sdAbs. The acquisition of polybasic sites uniquely by SARS-CoV2 S protein in comparisons to the other members of *coronaviridae* is considered to enhance the virus infectivity^30^. The proteolytic enzymes are abundantly present at infected tissue sites as well as in the membrane of susceptible cells and can process S protein to generate S1 and S2 subunits^31,32^. Such events facilitate the virus entry mediated by surface receptors such as ACE2 in the susceptible cells^31,33,34^. As polybasic sites serve as one of the virus virulence determinants for SARS-CoV2, we considered this as a potential target for selecting sdAbs. Such reagents could not only be valuable for neutralizing the virus infectivity but also help deciphering events in internalization and trafficking^35^. SARS-CoV2 generates variants and the efficiency of such processes increase under the influence of selective immune pressure^36^. Many mutations have been mapped in the RBD of S protein rendering the virus neutralization by antibodies less efficient^37^. Therefore, additional targets need to be identified and monoclonal antibodies such as the anti-CS sdAb could be considered as a potent therapeutic strategy. We also speculate that such reagents could potentially be useful in detecting infected cells by flow cytometry and understanding the biogenesis, multiplication and trafficking events of SARS-CoV2 within the susceptible cells.

## Materials and Methods

### Prediction of B cell epitopes

Linear B cell epitopes were predicted by IEDB database using BepiPred 2.0 server. The amino acid sequence of spike glycoprotein (PDB ID: 6VSB) was used as the input sequence^38^. A threshold value of 0.50 was used for predicting peptides to define the specificity and sensitivity of immunogenic epitopes. The epitopes were selected based on the individual score and the target region of the spike protein. The prediction epitopes were commercially synthesised from GL Biochem Ltd. (Shanghai, China) and were used for biopanning sdAb against SARS-CoV2 S protein.

### Generation of helper phage

Competent TG1 bacterial cells were cultured in glucose supplemented 2xYT medium until an OD_600_ value of 0.4 was reached. TG1 cells were then infected with helper phage M13K07 under static conditions at 30°C for 40 mins. Thereafter, the medium was supplemented with kanamycin (50μg/ml) and grown overnight for the multiplication of phages. Bacterial cells were then pelleted at low temperature to collect supernatant. The supernatant was then precipitated using 20% polyethylene glycol (PEG) and 0.5M NaCl to obtain helper phages, used for infecting recombinant TG1 cells harbouring VHH sequences.

### Recombinant phage preparation and Bio panning

Helper phage particles expressing VHH that were then used for bio-panning as described earlier^22^. Synthetic peptides predicted to be immunogenic for B cells were coated onto ELISA plates (50μg/ml/well) at 4°C overnight followed by blocking the wells with 4% BSA for 2hrs at room temperature (RT). ELISA plates were then washed three times with freshly prepared phosphate buffer saline with 1% Tween-20 (PBST). Subsequently, 10^12^ recombinant phages/well were added to the plate followed by incubation for 3 hrs at RT. Unbound phages were removed by extensive washings (25 times) with a freshly prepared PBST. The bound phages were then eluted using freshly prepared alkaline triethylamine acetate (TEA) buffer. The eluted phages were further enriched to enhance the affinity of peptide specific VHH by performing second round of bio-panning. The eluted phages were then used for infecting TG1 bacterial cells.

### Cloning and expression of sdAbs

Multiple bacterial colonies obtained were screened by colony PCR using VHH specific primers as described earlier^22^. The positive clones were selected to isolate phagemids. The retrieved VHH sequences were further sub-cloned into a modified pYBNT Vector containing a T7 promoter for bacterial expression^22^. Out of the multiple colonies obtained, two clones were further processed for producing the sdAbs against each of the target peptides from the RBD and the CS of SARS-CoV2 S protein. Clones were sequenced, expressed, purified, and characterized further. A primary bacterial culture of desired clones was propagated overnight in LB media supplemented with ampicillin (100μg/ml) at 37°C followed by scaling up as one liter culture in shaking flask until an OD600 of 0.4-0.6 was reached. The cultures were then induced with 1mM of IPTG for the induction of recombinant protein at 37°C in a shaking flask for 4 hours. Thereafter the cultures were pelleted down by centrifugation at 8000rpm for 10 minutes at 4°C.

### Purification of sdAb from inclusion bodies

To purify the recombinant protein from inclusion bodies, the pellet was first resuspended in lysis buffer containing 100mM Tris base and 10mM EDTA. The bacterial suspension was sonicated on the ice at an amplitude of 40 with 8 cycles with one-minute pulse on and one-minute off. Subsequently, the cells were centrifuged at 8000rpm for 10 minutes at 4°C to obtain the pellets which were then washed twice with wash buffer A (100mM Tris base, 10Mm EDTA, 1M NaCl, pH 8.0) and once with wash buffer B (100mM Tris Base, 10Mm EDTA, 1% v/v Triton 100; pH 8.0). The pellets were finally resuspended in denaturation buffer containing 100mM NaH_2_PO_4_, 10mM Tris-HCl, 8mM urea, pH 8.0 and kept at rotation for 18-20 hours at 4°C. The denatured fractions were centrifuged at 5000 rpm at 4°C for 10 minutes to obtain the clear extract. The supernatant were subjected to Ni-NTA purification using His-trap columns pre-equilibrated with the denaturation buffer. The washing was done with 20mM imidazole containing denaturation buffer (pH 8.0). The bound product was eluted using 400mM imidazole in denaturation buffer (pH 7.8). The final protein yield was 20mg/liter. The protein thus obtained was mixed with an equal volume of guanidine solution (3M GuHCl, 10mM sodium acetate and 10mM EDTA, pH 4.2) and was refolded by a rapid dilution method using 100mM Tris, 1mM EDTA, 1mM GSH, 0.1mM GSSG, 400mM arginine as described earlier^39^. The refolded fraction was then subjected to size exclusion chromatography using an S200 Hiprep column and two dominant peaks were obtained. The peaks obtained were pooled separately and spectral analysis was done for both using circular dichroism.

### Measuring thermal and pH stability of sdAb

To measure the effects of pH on the structural integrity of purified sdAbs, the preparations were incubated in the buffers with varied pH values ranging from 2 to 13 for 10 minutes and CD spectral analysis was performed. Similarly, the effect of temperature on purified sdAbs was analysed by performing thermal kinetics. The sdAbs preparations were subjected to heating at different temperatures ranging from 20°C to 70°C and then cooling from 70°C to 20°C to measure their denaturation and renaturation kinetics.

### Generation of SARS-CoV2 pseudovirus

Plasmids encoding SARS-CoV-2 S protein were a kind gift from Dr. McLellan of the University of Texas, Austin and were described earlier^23,40^. Third generation lentiviral packaging vector, pCMVR.74 and pMD2.G (VSV-G envelop vector) were also used. Plasmid encoding SARS-CoV2-spike protein, pCMV14-3X-FLAG-SARS-CoV-2S was from Addgene (#145780) and was described earlier^41^. pLenti-GFP (Core with 5’ and 3’ LTR) was also from Addgene (#17448) and the plasmids containing Tat1b and Rev1b (SARS-Related Coronavirus 2, Wuhan-Hu-1 Spike-Pseudotyped Lentiviral Kit, NR-52948) were also obtained from BEI resources and were described earlier^42^. The above mentioned 5 plasmids, Spike construct (6μg), pCMVR8.74 (9μg), pLentiGFP (10.8μg), Tat-containing plasmid (μg) and Rev plasmid (6μg) were mixed with 1:3 of PEI (1μg/ml)^43^ in 5ml of serum free DMEM/petri-plate, followed by instant mixingby vortexing for 40 seconds. Above mixture was kept at RT for 15 minutes and co-transfected in HEK239T cells (Human Embryonic Kidney). The transfected HEK293T cells were used for the generating replication incompetent pseudotyped lentivirus (LV) expressing SARS-CoV2 S protein labelled as LV(CoV2-S) and VSV-G protein labelled as LV(VSV-G). The supernatants were collected at 72hrs post transfection and centrifuged at 1000g to remove cell debris. The collected viral supernatant was concentrated with a solution, 1.2M NaCl and 40%(W/V) PEG-8000 on ice for 3-4 hrs followed by centrifugation at 3500g for 65 minutes. The precipitated virus pellets were dissolved in serum free medium, stored at 4°C and used within a few days-time. Modified Vero cells (Vero E6) originated from kidney epithelial cells of African green monkey were used for measuring the internalization using concentrated pseudoviruses. The cells were grown in 10% DMEM supplemented with 10x Penicillin/Streptomycin and maintained in a humidified CO_2_ incubator at 37°C. Cellular infections with pseudoviruses were performed in serum free medium and after 24 hrs the culture media were replaced with complete DMEM. The cells were analysed for GFP expression 72 hrs post-infection by fluorescence microscopy using Nikon eclipse Ti. The cells were detached from the culture plates using 1mM PBS-EDTA, GFP expression was measured by flow cytometry. Vero E6 cells infected with bald particles or the LV (VSV-G) pseudovirus particles served as a control for such experiments.

### Measuring neutralization of pseudoviruses by sdAbs

Different dilutions of anti-CS and anti-RBD sdAbs were used for measuring the internalization of LV(CoV2-S) pseudovirus particles by Vero E6 cells. For blocking experiments, we pre-incubated different concentrations of the sdAbs (1ng/ml, 10ng/ml, 100ng/ml, 500ng/ml, 1μg/ml, 5μg/ml and 10μg/ml) with 5×10^6^ of either LV (CoV2-S) or LV(VSV-G) on ice for 1 hr. Subsequently, the mix was added to 70-80% confluent Vero-E6 cells. After 24 hrs medium was replaced with a fresh complete medium. The supernatant collected from these cells were analyzed by SDS-PAGE and western blotting. At different time points post-infection, the cells were analysed for GFP expression by fluorescent microscope and flow cytometry (BD Accuri).

In order to measure the specificity of sdAbs, anti-CS and anti-RBD sdAbs were preincubated with different concentrations of their respective peptides for 1 hr on ice, followed by adding these mixtures to LV(CoV2-S) as described earlier. GFP expression was measured 72 hrs post-infection using fluorescence microscopy and flow cytometry. In additional experiments, the anti-CS antibody was added to trypsin-treated LV(CoV2-S) for 4 hrs at 37°C, followed by incubation with anti-CS sdAb (100ng/ml) for 1 hrs on ice. The above mixture was applied to Vero E6 cells and the infectivity was measured. Aliquots of the same sample was also for ELISA as well as western blotting to analyse to analyse the trypsin mediated cleavage of SARS-CoV2-S in the pseudovirus particles.

### Measuring the effect of specific sdAbs in fusogenic activity

HEK293T cells were transfected with different plasmids to make LV (CoV2-S) VLPs and Spike protein. After 72 hrs the generated pseudovirus particles were collected and the remaining transfected cells, HEK293T^+LV(CoV2-S)^, were scrapped and used for cell fusion assay. The expression of the S protein in HEK293T cells was measured by their surface staining with both the sdAbs using flow cytometry. HEK293T^+LV(CoV2-S)^ were incubated with different concentrations of anti-CS and anti-RBD sdAbs for 1hr on ice followed by their co-culture with Vero E6 cells for 6 hrs at 37°C in humidified CO_2_ incubator. The frequencies of fused cells were measured by fluorescence microscopy and flow cytometry^27^. The fused cells would increase in size, therefore a flow cytometric analysis was performed.

### Flow cytometry

After 72 hrs of transduction, Vero-E6 cells were treated with 1mM PBS-EDTA for 15 minutes at 37°C in CO_2_ incubator and the cells were removed from 96 well (flat bottom) plates by reverse pipetting. Cells were collected in 1.5ml micro-centrifuge tube, washed twice and acquired using flow cytometer (BD Accuri). The available data was analysed using flowJo X software (TreeStar)^44^.

### Fluorescent Microscopy

The cells were analysed for GFP expression 72 hrs post-infection by fluorescence microscopy using Nikon eclipse Ti and all images were taken at 10X magnification. Analysis and scaling of all the taken images were done using ImageJ software^44^.

### Scanning electron microscopy

Field Emission Scanning Electron Microscope (FESEM) was used to measure the surface topography of LV(CoV2-S) and LV(VSV-G). Pseudoviral particles were spread and dehydrated on glass slide overnight, followed by coating with gold nanoparticle for providing conductivity to the samples and images were acquired using JEOL JSM-7600F FESEM.

### Western Blotting

To determine the specificity and immune reactivity of the sdAbs with the SARS-CoV2 S protein purified from transfected HEK293T cells or the one displayed by the produced pseudoviruses, the polypeptides in the prepared samples were resolved using SDS-PAGE and after transferring to PVDF membrane were immunoblotted with the sdAb containing 6x(HIS)-tag or their biotinylated versions after blocking of the membrane with 5% skim milk. Anti-6x(HIS) anti-mouse antibody was used to recognize VHH (4A12E6: Invitrogen Rockford, USA) followed by probing with anti-mouse IgG raised in goat (A3562: 1ml Sigma, USA) and conjugated with alkaline phosphatase for the development of the blot. Culture supernatants and pseudoviruses expressing spike construct, LV(CoV2-S), Bald particles, LV(BALD), pseudoviruses expressing VSV-G protein, LV(VSV-G) were precipitated using 40% PEG −1.2M NaCl overnight on rotation at 4°C. The samples were centrifuged at 5000rpm for 40 min and washed with PBS and then incubated with 1x SDS sample buffer and 4x Laemmli sample buffer for 10mins at room temperature followed by heating for 15 min at 96°C. After resolving polypeptides using 12% SDS PAGE, the transferred onto PVDF membrane was performed. The membranes were then blocked with 5% skimmed milk for various durations. The blocked PVDF membrane was probed with the sdAb (having both 6x(HIS)-tag and biotin-tag) and then monoclonal anti-6x(HIS)-tag antibody was probed followed by detection with anti-mouse IgG conjugated with alkaline phosphatase (A3562: 1ml Sigma, USA) was used for developing the blots.

Since the spike construct contains the Flag tag, immunoblotting was performed to detect the presence of spike protein in supernatant collected 24 hours post transduction as well as in cell lysate of 72 hours post transduced Vero E6 cells using anti-flag mouse antibody (F1804-50UG, USA). A secondary antibody anti-mouse IgG raised in goat and conjugated with alkaline phosphatase was used for the development of the blot.

### Indirect ELISA and competitive ELISA

Peptide against which sdAb was biopanned was coated at a concentration of 50μg/ml overnight at 4°C. The following day the wells were washed with 0.05% PBST and were blocked in 5% BSA in PBST for 2hrs at RT followed by washing with PBST and incubation with sdAb in different dilutions for 1.5 hrs at RT. For detection of sdAb, anti-6x(HIS) antibody was incubated for 1 hr at RT followed by washing and incubation with anti-mouse antibody conjugated with alkaline phosphatase for 1.5 hrs. The plate was washed and developed with 100μl of pNpp substrate from Sigma Aldrich (1mg/ml) in glycine buffer. 50μl of stopping solution (3M NaOH) after the development of colour was added and absorbance was taken at 405nm.

### Biolayer interferometry (BLI)

BLI was performed to determine the binding kinetics of anti-CS and anti-RBD antibody with the purified spike protein produced from transfected HEK293T cells (S construct with 3x FLAG-tag) using BLltz System. For loading sdAbs with 6x(HIS)-tagged, Ni-NTA probes (ForteBio) were used. 200□l of 250μg/ml of both antibodies were loaded for 5 minutes and then washed with PBS to remove non-specific binding. Different concentrations of S protein were incubated for 5 minutes to measure the binding affinity with the immobilised anti-CS and anti-RBD antibodies, and their dissociation kinetics was measured in PBS. For reusing the probes re-charging was done by placing the sensor in 10mM glycine for 1 minutes and then in PBS for 5 minutes. Finally, the probes were placed in 10mM Nickel sulphate solution for 1 minute.

### Statistical analysis

One way ANOVA (and non-parametric or mixed) Dunnett’s multiple comparisons test were performed for all the group analysis. The results are presented as mean ± SEM. The p values are shown in the figures are represented as ⋆p ≤ 0.05, ⋆⋆p ≤ 0.01, or ⋆⋆⋆p ≤ 0.001. All statistical analysis were done using Graphpad prism 8.0.2(263).

**Figure 6.**
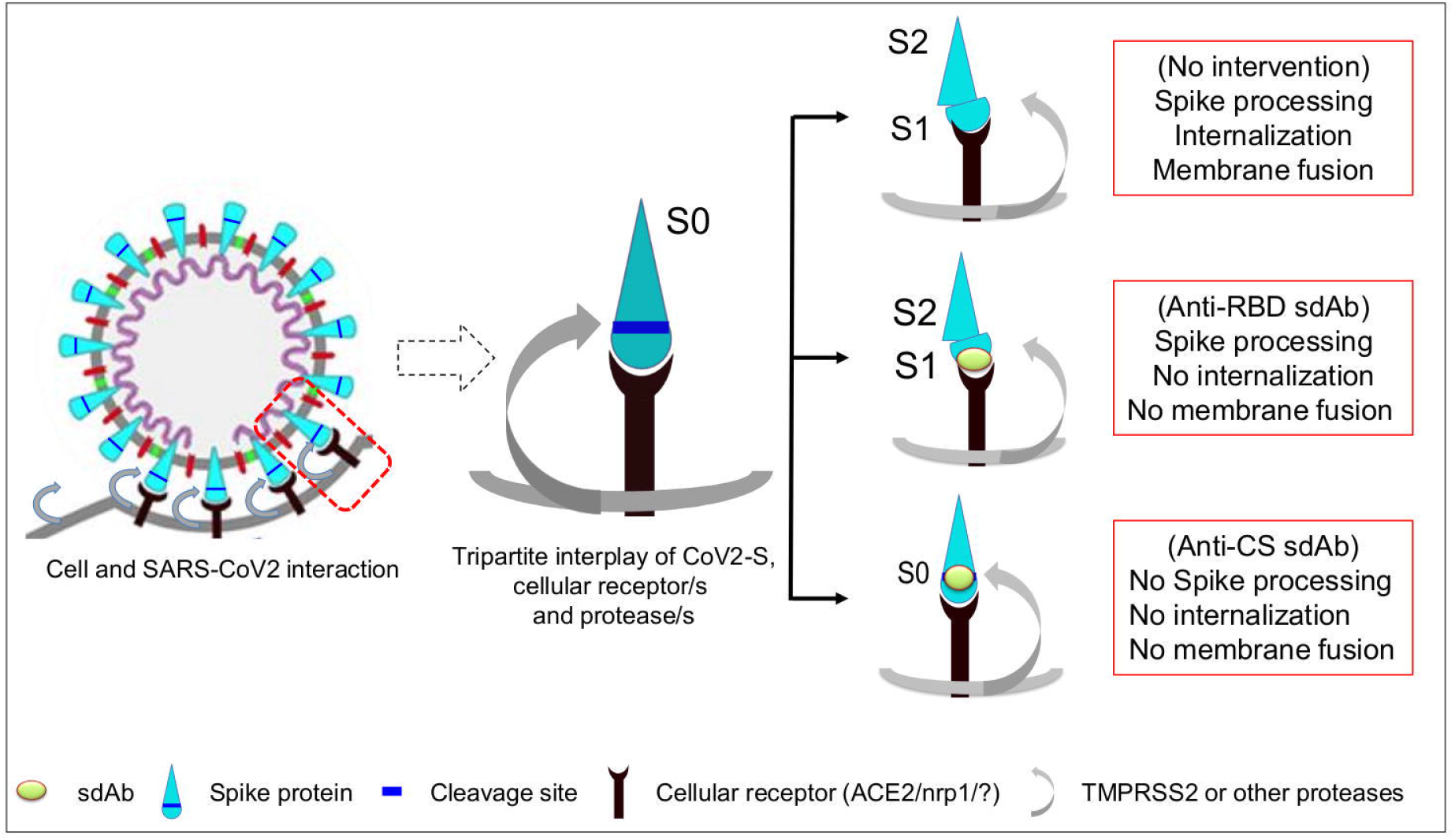
A cartoon shows the probable mechanism by which the anti-CS and anti-RBD sdAbs neutralise SARS-CoV2 infectivity. SARS-COV 2 S protein in its homotrimeric state interacts with its cellular receptor (ACE2/neuropilin 1/ some yet to be identified receptors) through RBD. This interaction is followed by host protease mediated cleavage of S protein (S0) to yield S1 and S2 fragments. Therefore, the targeting of RBD using sdAbs could block interaction with the cellular receptor to effect neutralization but the cleavage of the S protein might still occur. As the concentration of neutralizing antibody drops to suboptimal levels the virus could infect cells followed by its membrane fusion and the release of viral genome proceeds in subcellular compartments. Targeting the cleavage site however, could serve to increase the durability of anti-viral activity as the lack of S protein processing further reduces the chances of membrane fusion and internalisation of the virus.

## Acknowledgements

We thank Professor Kartik Chandran and Dr Rohit K Jangra of Albert Einstein college of Medicine, New York and Dr Jason McLellan of the University of Texas, Austin for sharing critical reagents for the work. The initial work for preparing the phage display library was done by Dr Manpreet Kaur. We also thank Dr Indranil Banerjee, Professor Anand K Bachhawat and Dr Anant Venkatesan for discussion and additional support.

