## Supplementary figures and images for "Targeting conserved viral virulence determinants by single domain antibodies to block SARS-CoV2 infectivity"

### FIgure S1

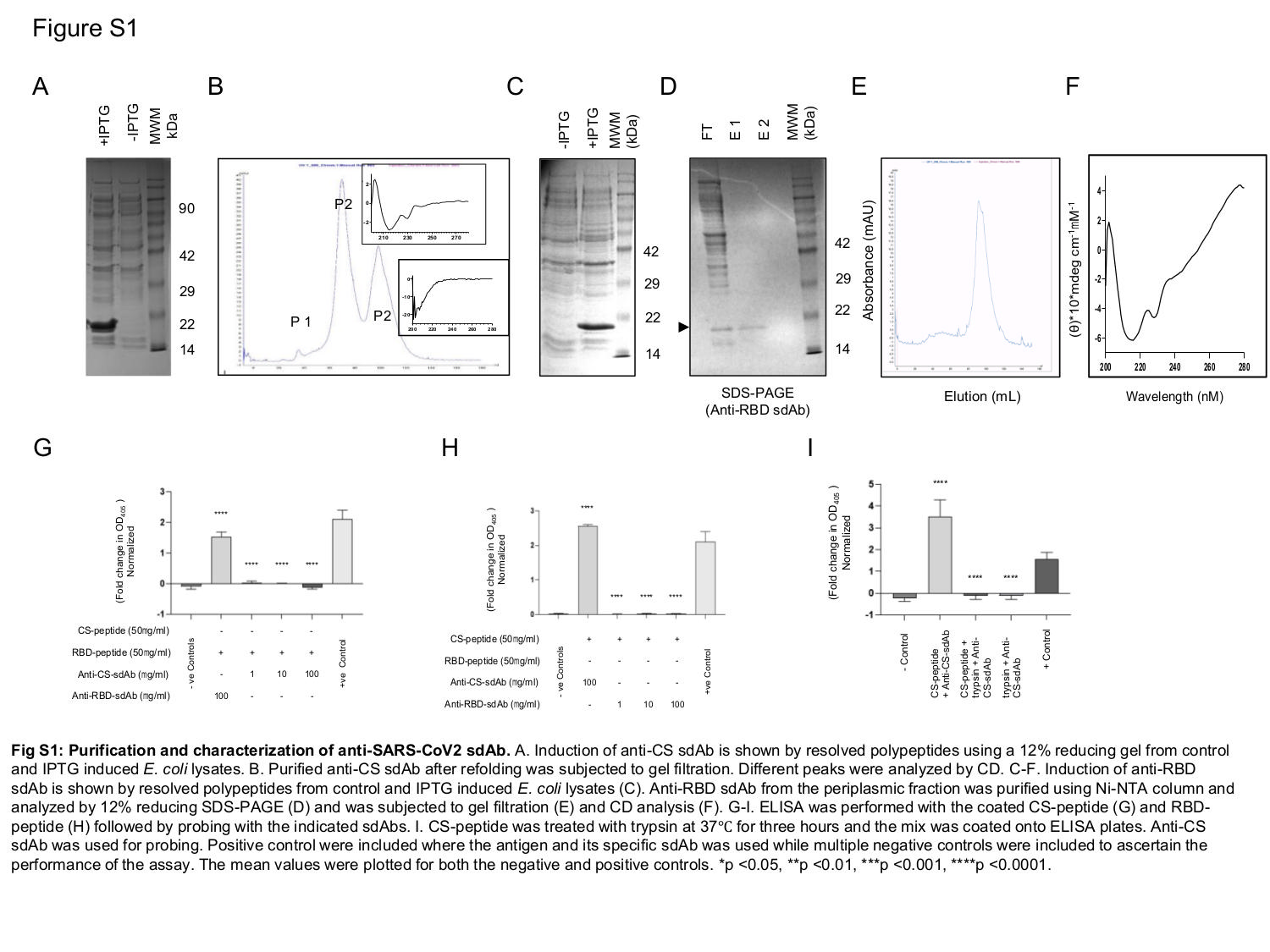

### FIgure S2

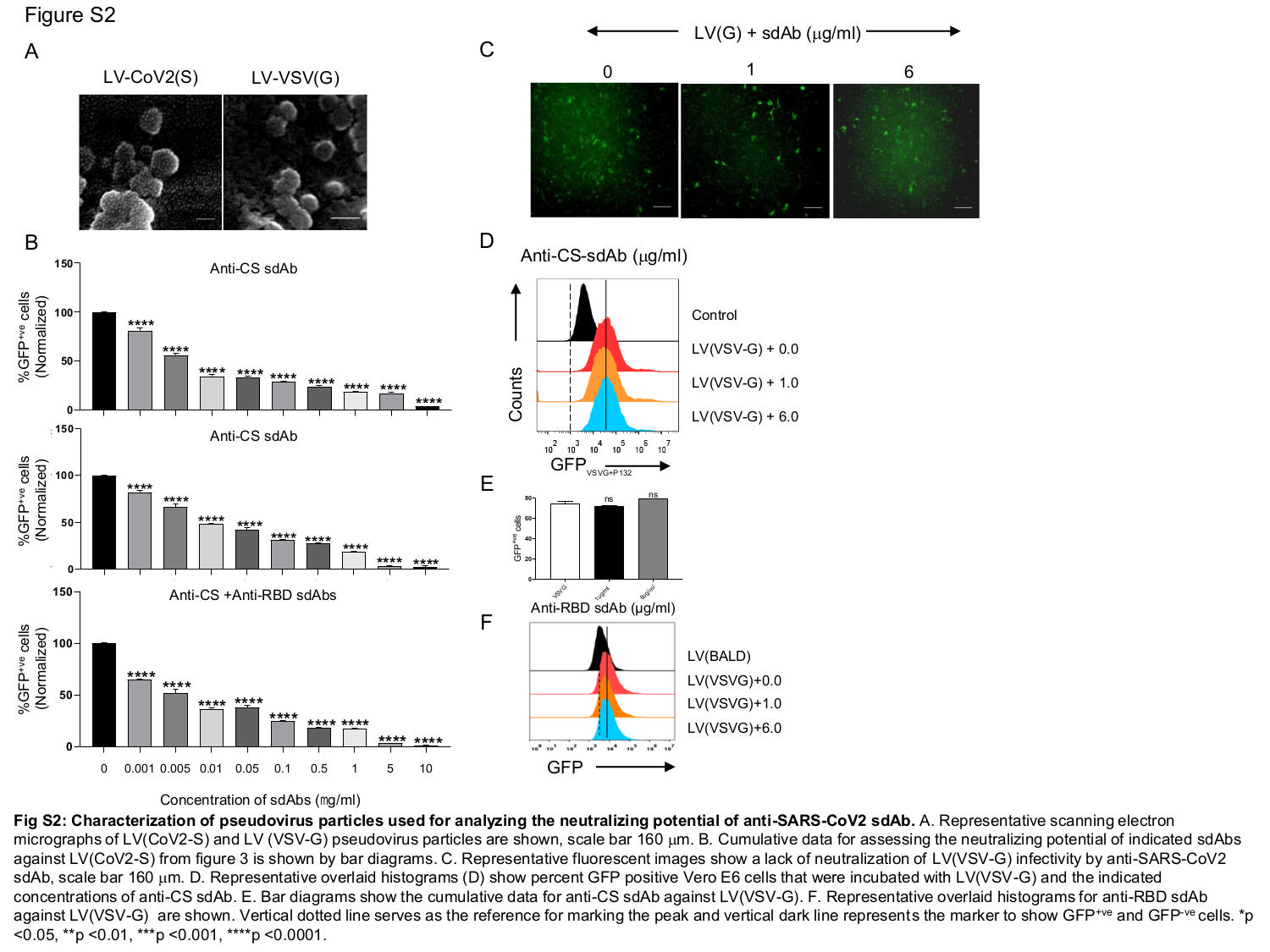

### FIgure S3

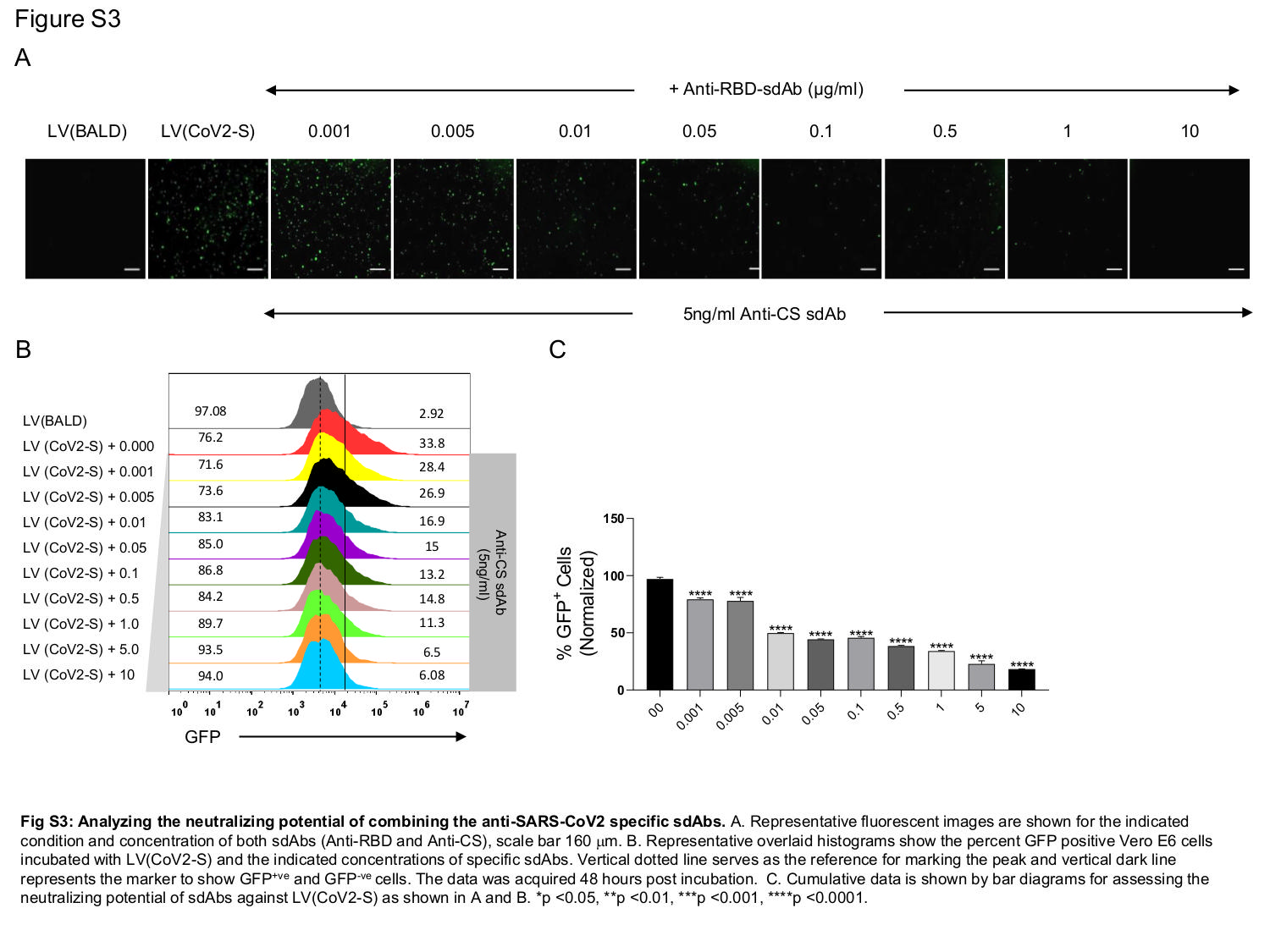

### FIgure S4

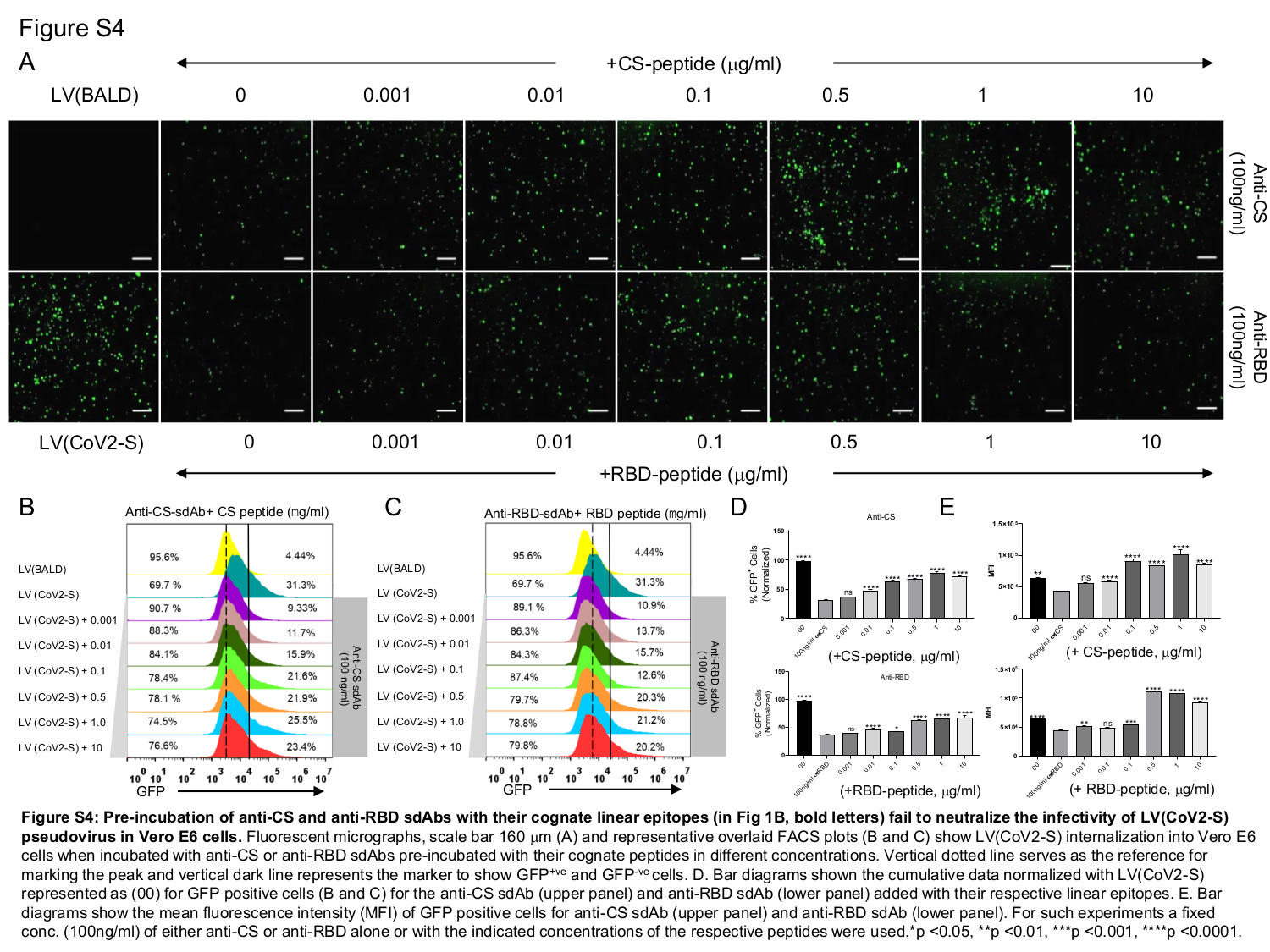

### FIgure S5

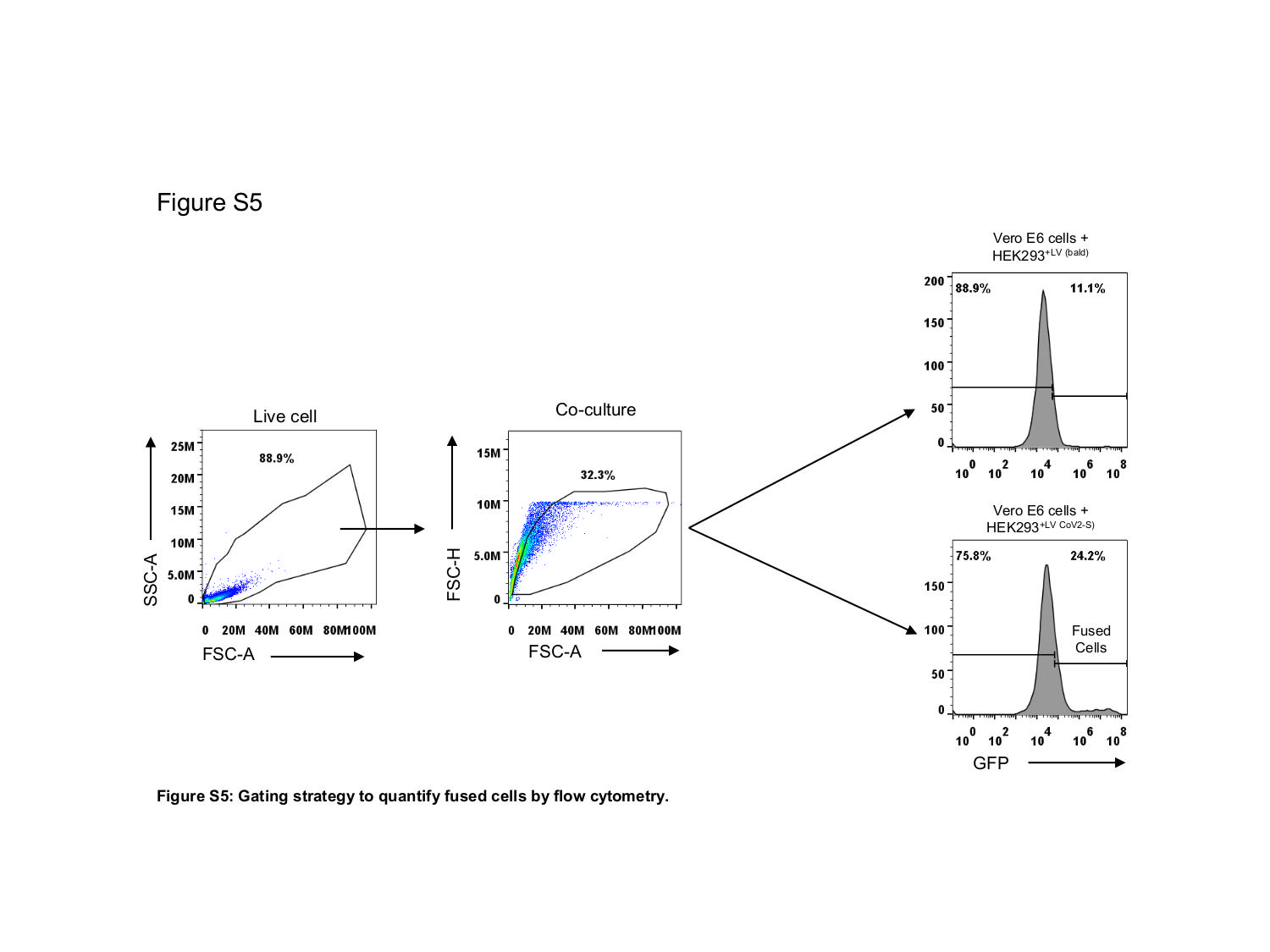

### FIgure S6

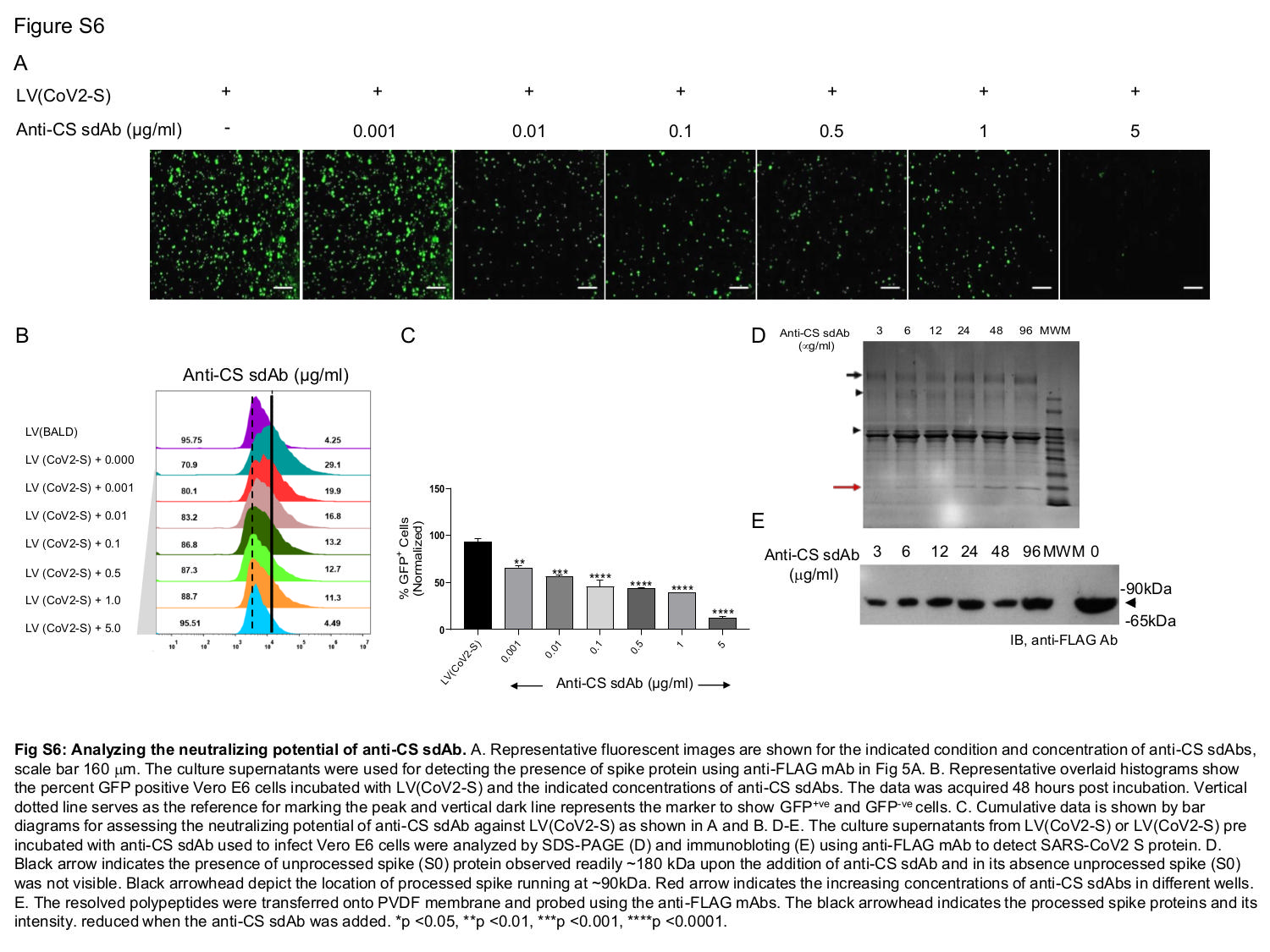

### FIgure S7

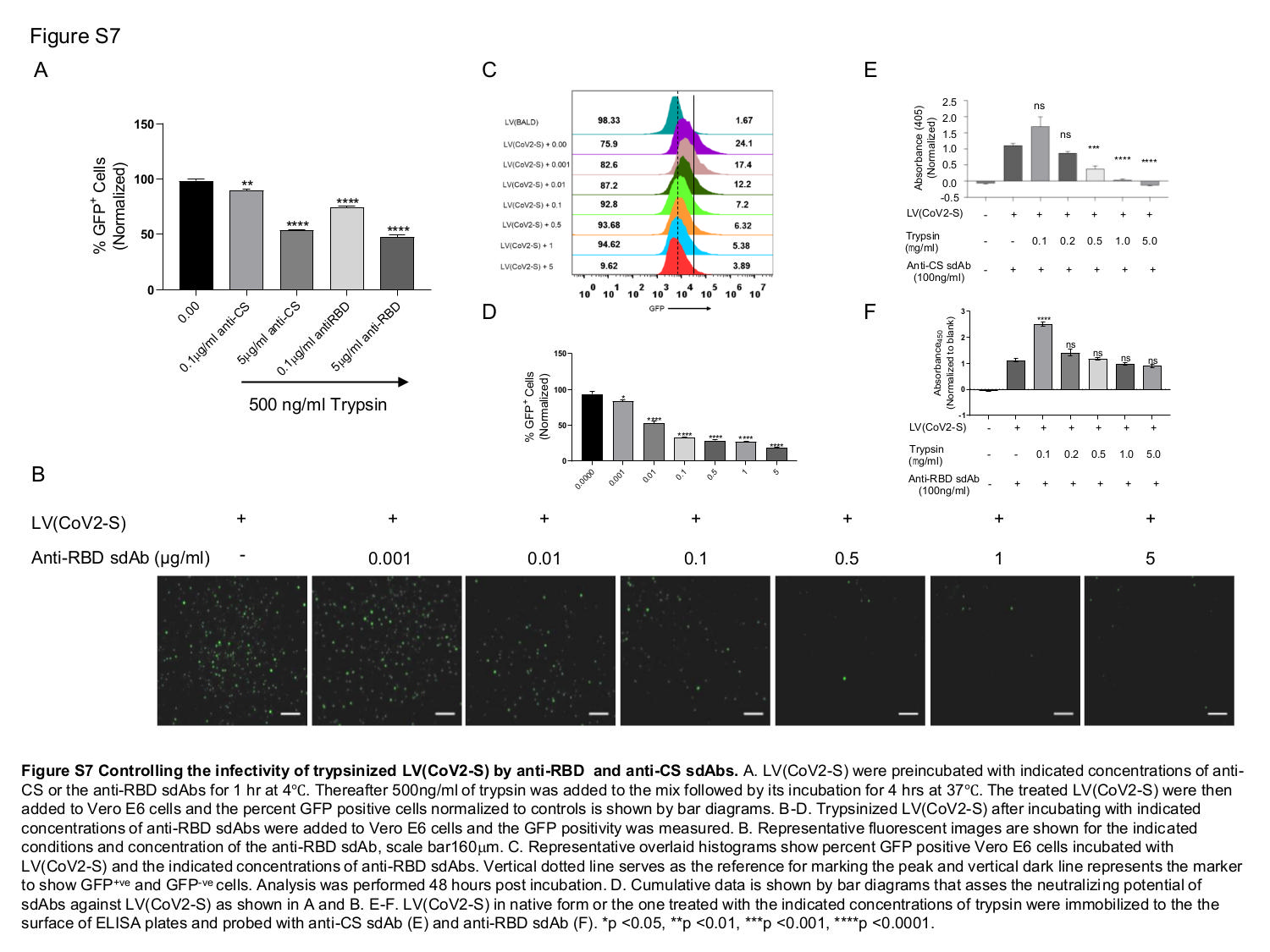
